# Transcriptome sequence reveals candidate genes involving in the post-harvest hardening of trifoliate yam *Dioscorea dumetorum*

**DOI:** 10.1101/2021.02.16.431375

**Authors:** Christian Siadjeu, Eike Mayland-Quellhorst, Sascha Laubinger, Dirk C. Albach

## Abstract

Storage ability of *D. dumetorum* is restricted by a severe phenomenon of post-harvest hardening which starts 72h after harvest and renders tubers inedible. Previous work has only focused on the biochemistry changes affecting the PHH on *D. dumetorum*. To the best of our knowledge nobody has identified candidate genes responsible for hardness on *D. dumetorum*. Here, transcriptome analysis of *D. dumetorum* tubers was performed, 4 months after emergence (4MAE), after harvest (AH), 3 days AH (3DAH) and 14 days AH (14DAH) on four accessions using RNA-Seq. In total between AH and 3DAH, 165, 199,128 and 61 differentially expressed genes (DEGs) were detected in Bayangam 2, Fonkouankem 1, Bangou 1 and Ibo sweet 3 respectively. Functional analysis of DEGs revealed that genes encoding for cellulose synthase A, xylan O-acetyltransferase chlorophyll a/b binding protein 1,2,3,4 and transcription factor MYBP were found predominantly and significantly up-regulated 3DAH, implying that genes were potentially involved in the post-harvest hardening. A hypothetical mechanism of this phenomenon and its regulation has been proposed. These findings provide the first comprehensive insights into genes expression in yam tubers after harvest and valuable information for molecular breeding against the post-harvest hardening. A hypothetical mechanism of this phenomenon and its regulation has been proposed. These findings provide the first comprehensive insights into genes expression in yam tubers after harvest and valuable information for molecular breeding against the post-harvest hardening.

## 1. Introduction

Yams constitute an important food crop for over 300 million people in the humid and subhumid tropics. Among the eight yam species commonly grown and consumed in West and Central Africa, *Dioscorea dumetorum* is the most nutritious [1]. Tubers of *D. dumetorum* are rich in protein (9.6%), well balanced in essential amino acids (chemical score of 0.94) and its starch is easily digestible [2]-[3]. *Dioscorea dumetorum* is not only used for human alimentation but also for pharmaceutical purposes. A bio-active compound, dioscoretine, has been identified in *D. dumetorum* [4], which is acceptable pharmaceutically and which can be used advantageously as a hypoglycemic agent in situations of acute stress. In Nigeria, the tuber is, therefore, used in treating diabetes [5].

Despite of these qualities, the storage ability of this yam species is restricted by severe post-harvest hardening (PHH) of the tubers, which begins within 24 h after harvest and renders them unsuitable for human consumption [1]. The post-harvest hardening of *D. dumetorum* is separated into a reversible component associated with the decrease of phytate and an irreversible component associated with the increase of total phenols [6]. The mechanism of post-harvest hardening is supposed to start with enzymatic hydrolyzation of phytate and subsequent migration of the released divalent cations to the cell wall where they cross-react with demethoxylated pectins in the middle lamella. This starts the lignification process in which the aromatic compounds accumulate on the surface of the cellular wall reacting as precursors for the lignification [7].

Whereas physiological changes associated with hardening of yam tubers are now reasonably well understood, we lack the knowledge of how to overcome hardening. Naturally occurring genotypes lacking post-harvest hardening [8] which offers a chance to understand the genetic basis of hardening. The next step has been to understand the genetic background of this genotype and its relationship to other genotypes, which has been conducted using GBS (Illumina-based genotyping-by-sequencing; [9]. Further insights have been gained by sequencing and analyzing the genome of the genotype Ibo sweet 3 [10].

Here, we analyze the transcriptome of this genotype Ibo sweet 3 and related genotypes to identify genes involved in the post-harvest hardening phenomenon. The study of the transcriptome examines the abundance of mRNAs in a given cell population and usually includes some information on the concentration of each RNA molecule, as a factor of the number of reads sequenced, in addition to the molecular identities. Unlike the genome, which is roughly fixed for a given cell line when neglecting mutations, the transcriptome varies from organ to organ, during development and based on external environmental conditions. In particular, transcriptome analysis by RNA-seq enables identification of genes that have differential expression in response to environmental changes or developmental stage and mapping genomic diversity in non-model organisms [11]. Differential gene expression analysis under different conditions has, therefore, allowed an enormously increased insight into the responses of plants to external and internal factors and into the regulation of different biological processes. High-throughput sequencing technologies allow an almost exhaustive survey of the transcriptome, even in species with no available genome sequence [12]. Indeed, transcriptome analysis based on high-throughput sequencing technology has been applied to investigate gene expression of hardening in carrot [13]. In yam, it helped elucidate flavonoid biosynthesis regulation of *D. alata* tubers [14].

A lack of availability of next generation ‘–omics’ resources and information had hindered application of molecular breeding in yam [15], which has recently been overcome by the publication of two genome sequences in the genus [10]-[16]. Here, we report the first transcriptomic study of *D. dumetorum* and the first to evaluate the influence of genes on the post-harvest hardening phenomenon in a monocot tuber using transcriptomics. We aim to close this gap by identifying candidate genes involved in the post-harvest hardening phenomenon of *D. dumetorum* to facilitate breeding non-hardening varieties of *D. dumetorum*.

## 2. Results

### 2.1. Descriptive statistics of RNA-Seq data

After trimming, 943,323,048 paired-end raw reads (150-bp in length) were generated. for 48 samples (Supplementary S1). Among these, 242.7, 224.6, 233.9 and 242,1 million reads were belonged to Bangou1, Bayangam2, Fonkouakem1 and Ibosweet3. On average, 90% of all the clean reads were aligned to reference genome. Furthermore, 56 % (on average) of those reads were uniquely mapped to the reference genome sequence. A PCA plot of the normalized read counts of all samples is depicted in Figure 1. The first two principal components (PCs) explained 69% of the variability among samples. Four months after emergence (4MAE) was distantly clustered from After Harvest (AH) and later on after harvest. No clear separation was observed between AH and later on after harvest (3DAH, 14DAH). However, taking into account accession specificity, AH is distantly grouped from 3DAH and 14DAH. This finding indicated a difference between transcriptome expressions of accessions after harvest. One biological replicate of each accession at a specific time point did not cluster with others likely due to individual variability between plants.

**Figure 1.**
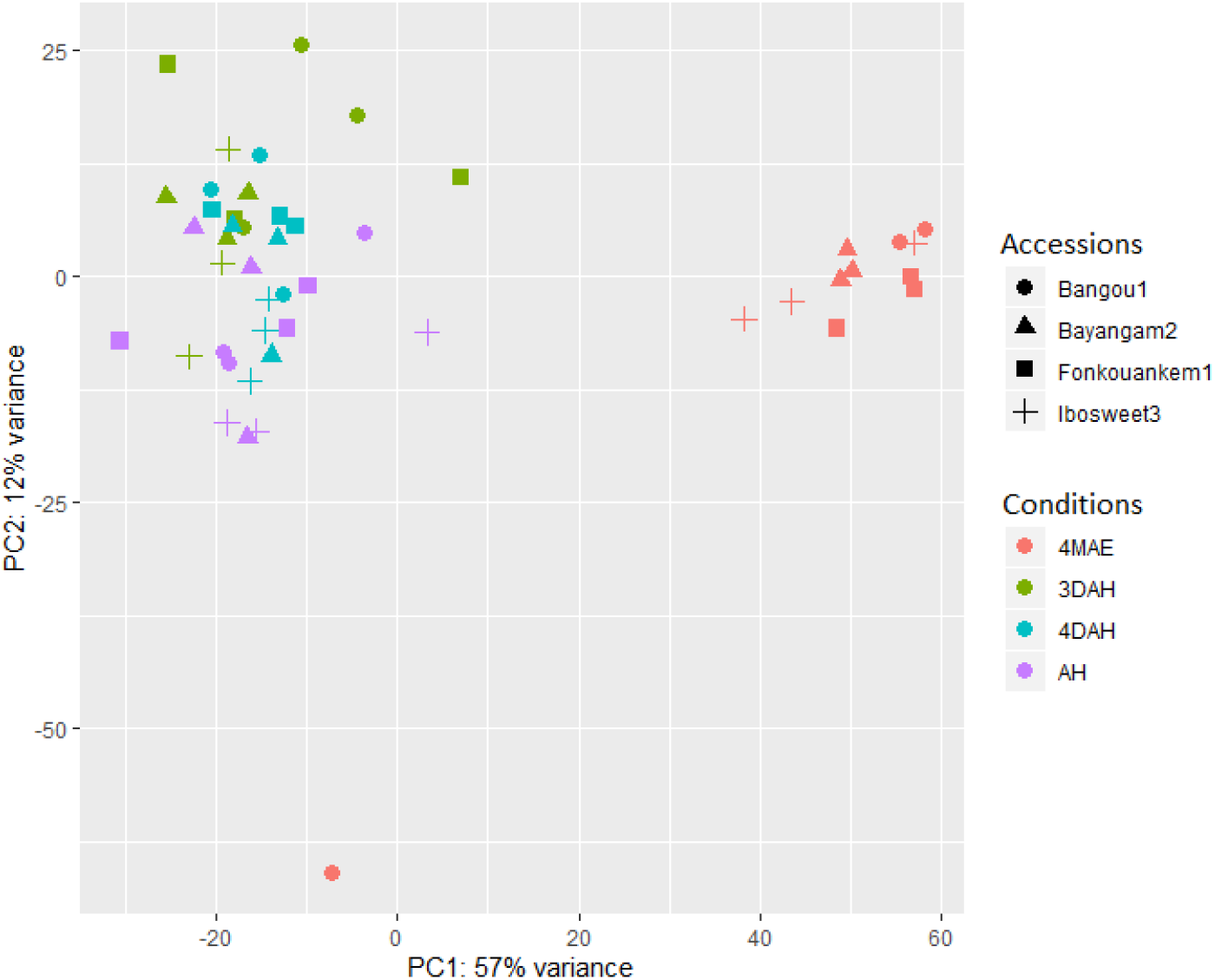
PCA plot of normalized count using VSD.

### 2.2. Differential expression analysis

Two well established statistical analysis methods to assess differentially expressed genes based on read counts (edgeR and DESeq2). We used two strategies to determine DEG on *D. dumetorum* after harvest: STAR_DESeq2, and STAR_edgeR. The design model for DE analysis was ~ Accession + Conditions + Accession:Conditions. We carried out multiple comparison at the accession, conditions and interaction accession*conditions levels. The approach STAR_DESeq2 yielded the highest number of DEG (Figure 2) and the results were selected for downstream analysis. Pairwise comparisons (4MAE vs. AH, 3DAH vs. AH, 14DAH vs. AH, 14DAH vs. 3DAH) of gene expression among the four accessions were performed (Figure 2). However, since the post-harvest hardening on *D. dumetorum* tubers occurs after harvest, results of gene expression were focused after harvest. A decrease of up-regulated DEGs and an increase of down-regulated DEGs were noticed among the 3 accessions that do harden from harvest to 14DAH (Figure 2). The accession that does not harden depicted a different pattern. Comparing 3DAH vs. AH, 165, 199,128 and 61 significantly DEGs were detected in Bayangam 2, Fonkouankem 1, Bangou 1 and Ibo sweet 3, respectively. Amongst these, 120, 112, 83 and 16 were up-regulated in Bayangam, Fonkouankem, Bangou1 and Ibo sweet3 respectively. For 14DAH vs. AH 162, 201, 161, and 46 significantly DEGs were obtained in Bayangam 2, Bangou 1, Fonkouankem 1 and Ibo sweet 3 respectively. Among which, 126, 83, 47, and 13 were up-regulated DEGs in Bayangam, Bangou1, Fonkouankem and Ibo sweet 3, respectively. In total, the highest number of significantly up-regulated DEGs were detected in Bayangam 2 and the lowest in Ibo sweet 3. A mixture analysis of all accessions that do harden irrespective of accession was performed (Supplementary S2). Pairwise comparisons of gene expression among the three stages (AH, 3DAH and 14DAH) detected 59, 40 and 13 up-regulated DEGs between 3DAH vs. AH, 14DAH vs. AH and 14DAH vs. 3DAH respectively. Whereas, 14, 36, and 56 down-regulated DEGs were obtained between 3DAH vs. AH, 14DAH vs. AH and 14DAH vs. 3DAH respectively.

**Figure 2.**
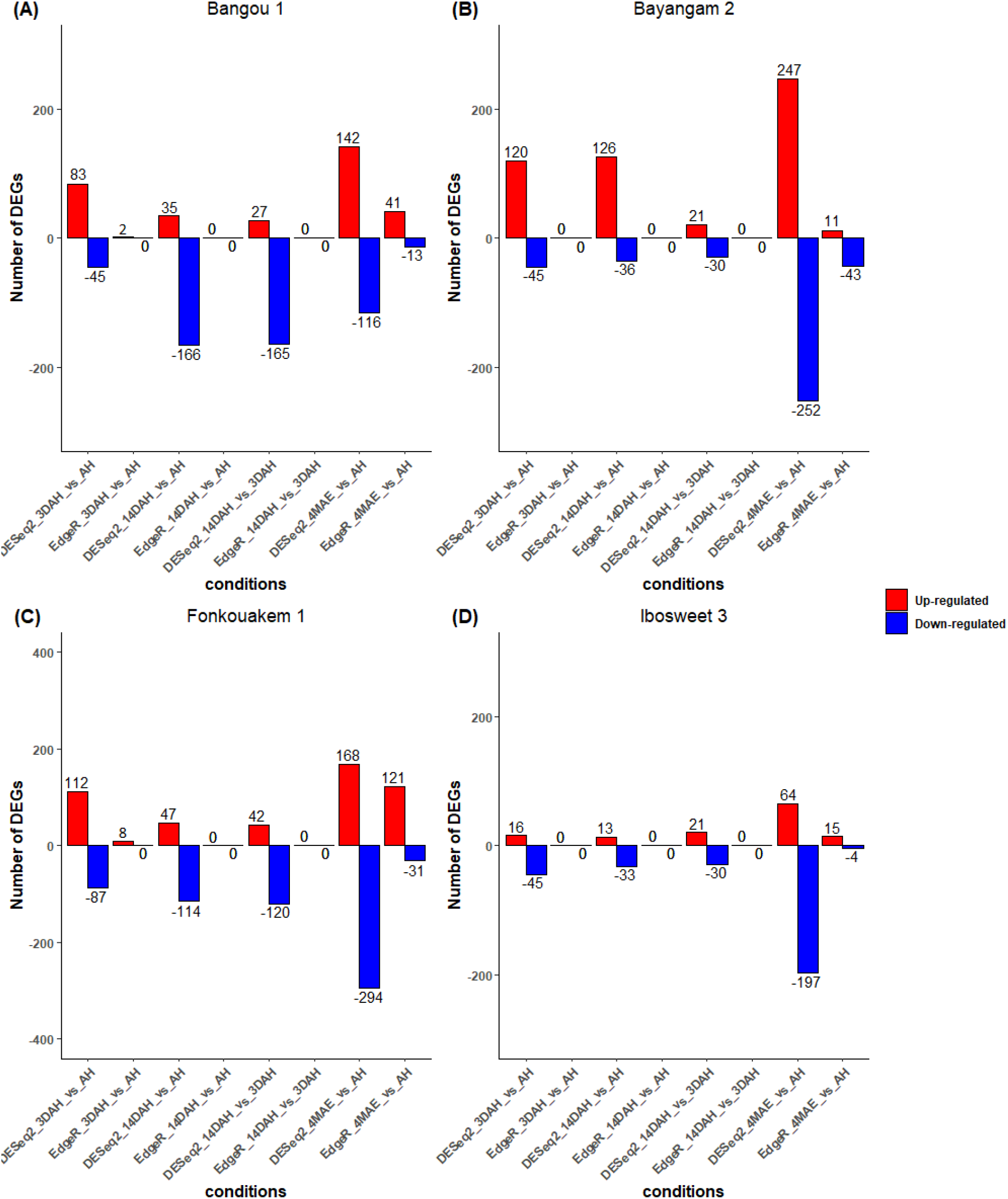
The number of DEGs based on the comparison of DESeq2 and EdgeR 4MAE and after harvest. (A), Bangou, (B), Bayangam (C), Fonkouankem, (D) Ibo sweet 3 (non-hardening accession). Blue represents down-regulated transcripts, and red represents up-regulated transcripts.

In order to understand the difference between Ibo sweet 3 and the other accessions, a multiple pairwise comparison (Bayangam vs. Ibo sweet 3, Bangou 1 vs. Ibo sweet 3, Fonkouankem vs. Ibo sweet 3) after harvest (3DAH vs. AH, 14DAH vs. AH) was carried out (Figure 3). After harvest to 3DAH (3DAH vs. AH), 111, 111 and 80 significantly DEGs were acquired comparing Bayangam vs. Ibosweet3, Fonkouankem vs. Ibo sweet 3 and Bangou1 vs. Ibo sweet 3 respectively. Amongst these, 101, 80 and 62 were up-regulated DEGs in Bayangam vs. Ibo sweet 3, Fonkouankem vs. Ibo sweet 3 and Bangou 1 vs. Ibo sweet 3 respectively. For 14DAH vs. AH, 88, 85 and 91 significantly DEGs were detected comparing Bayangam vs. Ibosweet3, Fonkouankem vs. Ibo sweet 3 and Bangou1 vs. Ibo sweet 3 respectively. Among which, 80, 30 and 22 were up-regulated in Bayangam vs. Ibo sweet 3, Fonkouankem vs. Ibo sweet 3 and Bangou 1 vs. Ibo sweet 3 respectively.

**Figure 3.**
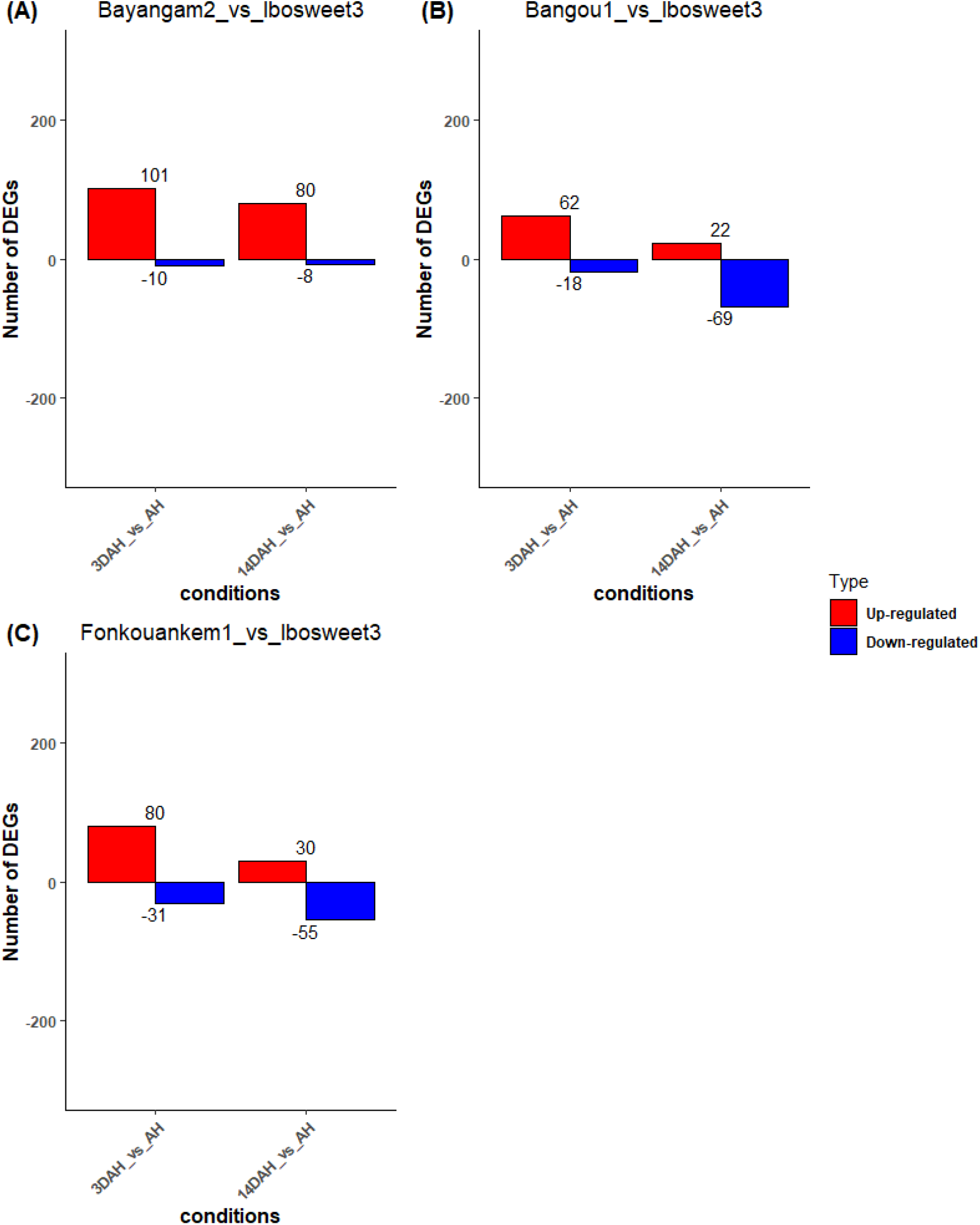
The number of DEGs based on the comparison between Ibo sweet 3 and other accessions after harvest. (A) Ibo sweet 3 vs. Bayangam 2, (A) Ibo sweet 3 vs. Bangou 1, (C) Ibo sweet 3 vs. Fonkouankem 1. Blue represents down-regulated transcripts, and red represents up-regulated transcripts.

### 2.2. GO enrichment and functional classification of DEGs with KEGG and Mapman

For better comprehension of the post-harvest hardening phenomenon, GO term annotation and enrichment was performed on up-regulated DEGs resulted from pairwise comparisons (3DAH vs. AH, 14DAH vs. AH) of all the three accessions that do harden (Figure 4 A). Compared with 3DAH and AH, out of the 59 up-regulated DEGs, 38 were significantly annotated in 43 GO terms, most of which were involved in biological processes related to cellular process, response to stimulus and metabolic process. Likewise, for 14 DAH vs. AH, 23 up-regulated genes (out of 40) were significantly enriched regarding biological processes in relation to cellular process, response to stimulus and metabolic process (Figure 4 B). Individual analysis of each accession separately revealed that cellular process, metabolic process, response to stimulus and response to stress belong to the top 10 of the mostly enriched GO term 3DAH and 14DAH for biological process (Figure 4 C, D).

**Figure 4.**
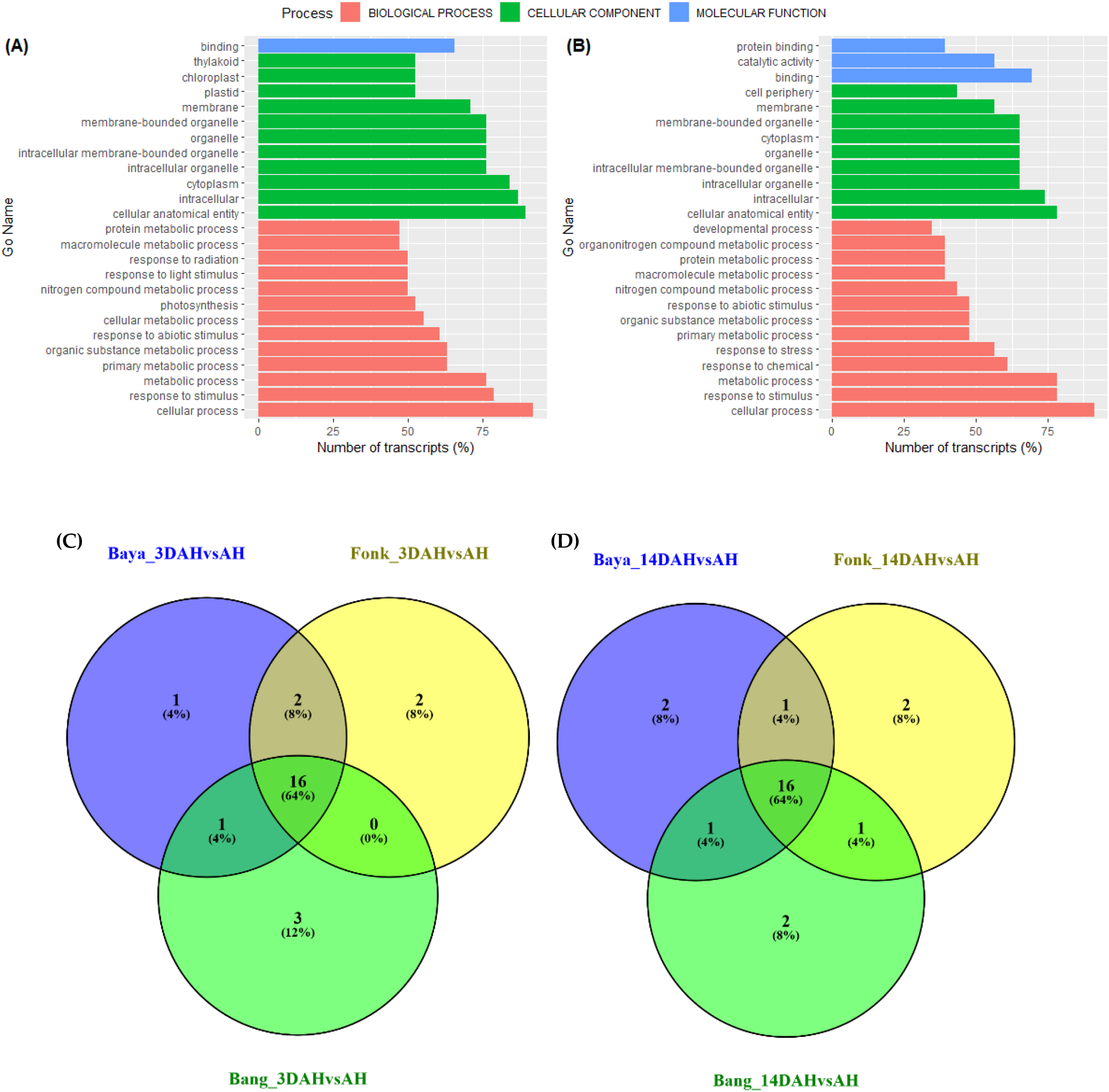
Functional annotation of the top 20 up-regulated enriched GO pathways of D. dumetorum tubers after harvest. (A) and (B) combined analysis of 3 hardened accessions 3 DAH and 14DAH respectively. (C) and (D) enrichment of each hardened accessions independently 3 DAH and 14DAH respectively. Blue bar represents molecular process, green bar represents cellular component, and red bar represents biological process.

Pathway-based analysis with KEGG revealed that metabolic pathway (Ko01100) was the most enriched with 7 and 6 up-regulated transcripts followed by biosynthesis of secondary metabolites (Ko01110) with 3 and 1 up-regulated transcripts 3DAH and 14DAH respectively (Figure 5 A, B). Based on MapMan photosynthesis pathway (Bin 1, 23 genes) and RNA biosynthesis (Bin 15, 8 genes) were the most enriched 3DAH. Likewise, 14 DAH, photosynthesis (6 genes) and RNA biosynthesis (6 genes) were the most enriched (Figure 4 A, B).

**Figure 5.**
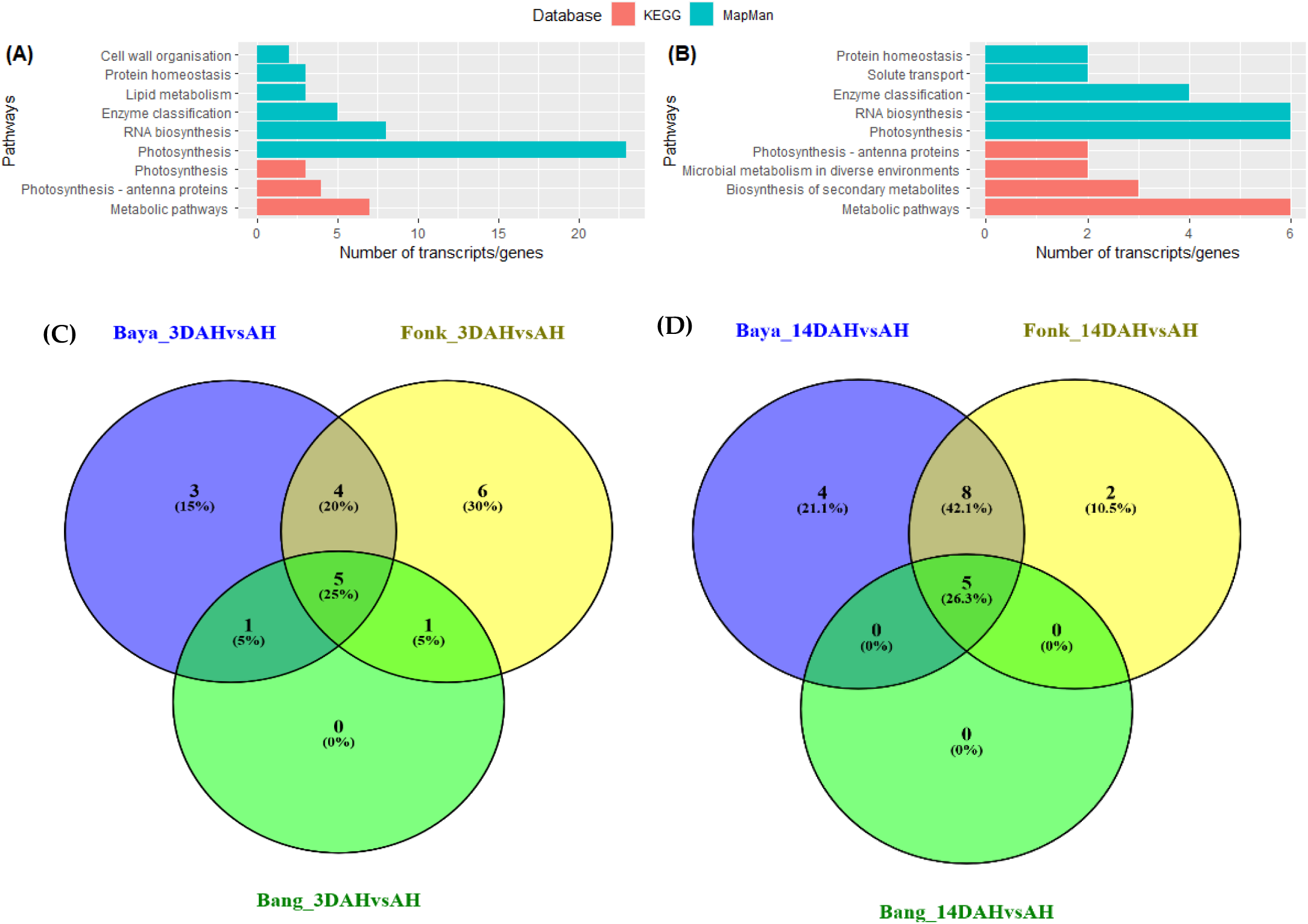
Functional classification of DEG after harvest. (A) and (B) the most enriched pathways of the combined analysis of 3 hardened accessions 3 DAH and 14DAH respectively. (C) and (D) the most enriched pathways of each hardened accessions 3 DAH and 14DAH respectively. Green bar represents pathway annotation with MapMan database, and red bar represents pathway annotation with KEGG database.

Pathway-based analysis with KEGG revealed that metabolic pathway (Ko01100) was the most enriched with 7 and 6 up-regulated transcripts followed by biosynthesis of secondary metabolites (Ko01110) with 3 and 1 up-regulated transcripts 3DAH and 14DAH respectively (Figure 5 A, B). Based on MapMan photosynthesis pathway (Bin 1, 23 genes) and RNA biosynthesis (Bin 15, 8 genes) were the most enriched 3DAH. Likewise, 14 DAH, photosynthesis (6 genes) and RNA biosynthesis (6 genes) were the most enriched (Figure 4 A, B).

### 2.3. Cluster expression analysis

Clustering gene expression of DEG 3DAH was assessed to identify groups of genes that are co-up-regulated (Figure 6). Two groups were identified amongst the genes differentially expressed 3DAH. One of the two clusters depicted a high peak 4MAE and then decreased AH and slightly increased 3DAH and 14DAH with an expression under zero except for the accession Fonkouankem. This group corresponds to cluster 1 for Bangou and Fonkouankem and cluster 2 for Bayangam and the mixture of the 3 accessions (Figure 6 A, B, C, D). Conversely, for the other cluster, the expression was down 4MAE and AH, and sharply increased 3DAH and then decreased 14DAH. This latter one showing the highest peak 3DAH is a group of genes that co-expression and could be involved in the post-harvest hardening. It corresponds to cluster 2 for Bangou and Fonkouankem and cluster 1 for Bayangam and the mixture of the 3 accessions. Therefore, functional annotation of genes of this group were further investigated.

**Figure 6.**
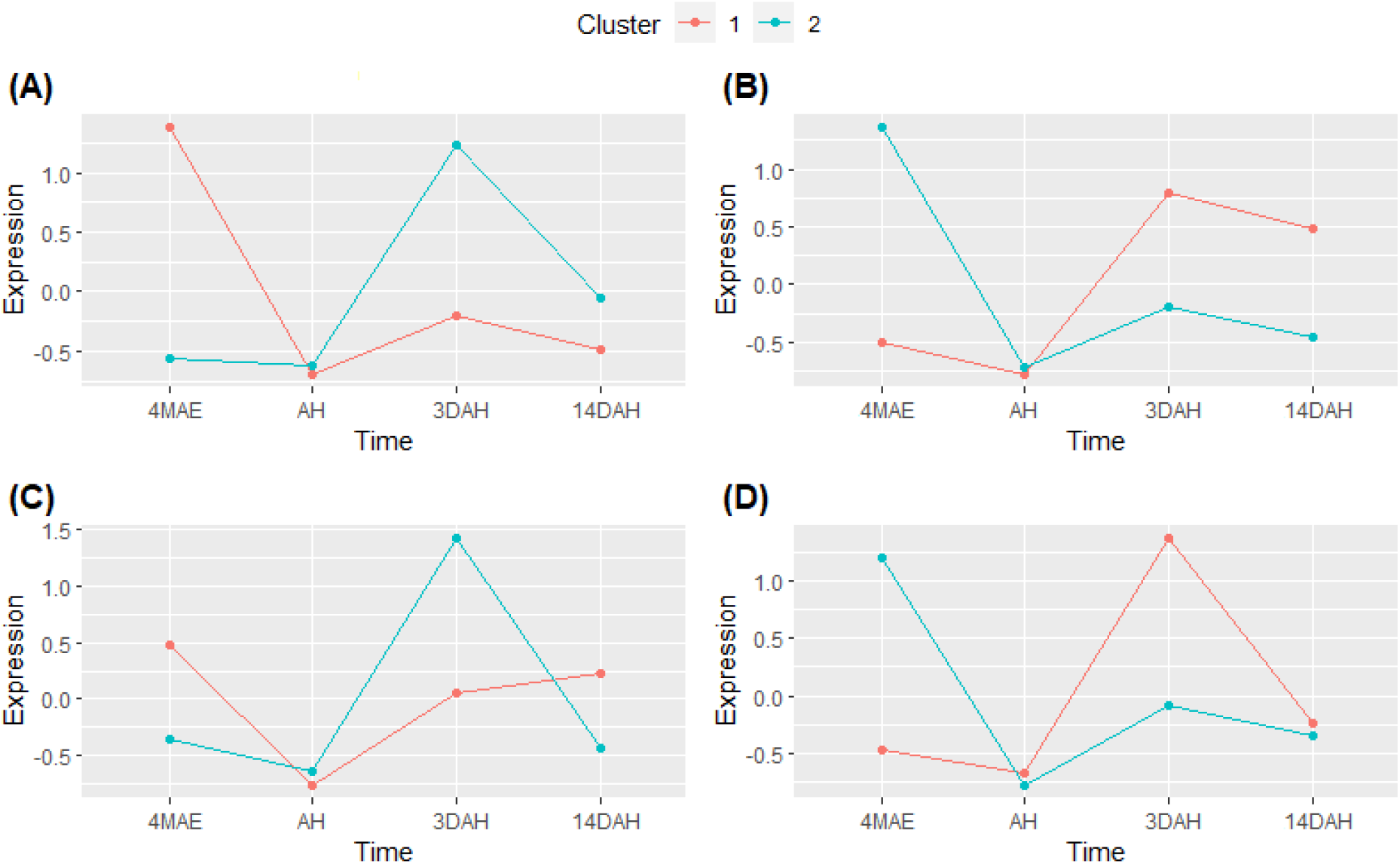
Cluster analysis of DEGs 3DAH among the different sampling time 4MAE and after harvest. (A), Bangou, (B), Bayangam (C), Fonkouankem, (D) combined analysis of the 3 hardened accessions.

The top 3 accumulated pathways in the cluster 2 were photosynthesis (20 contigs) followed by solute transport (2 contigs) and cell wall organization (1 contig) in Bangou (Supplementary S3). For Bayangam the top 3 where protein modification (8 contigs) followed by RNA biosynthesis (7 contigs) and phytohormone action (7 contigs). However, it is worth to outline that cell wall organization (4 contigs) and secondary metabolism (3 contigs) were as well accumulated. On the contrary in Fonkouankem cell wall organization (19 contigs) was the most enriched pathway followed by RNA biosynthesis (8 contigs) and photosynthesis, secondary metabolism, protein homeostasis, cytoskeleton organization and solute transport with 4 contigs each of them. The mixture of all those accessions showed that photosynthesis was the most accumulated pathway (21 contigs) followed by protein homeostasis, lipid metabolism with 3 contigs each of them and cell wall organization with 2 contigs. In sum, genes encoding for photosynthesis, cell wall organization, protein modification and RNA biosynthesis genes and secondary metabolism are co-up-regulated after harvest and likely involved in the post-harvest hardening on *D. dumetorum* tubers.

### 2.4. Comprehensive analysis of expression of genes potentially involved in post-harvest hardening

We opted for investigation of genes differentially expressed 3 DAH on the accession Fonkouankem due to its high amount of up-regulated genes associated with cell organization and the combining analysis of all three accessions together. In the cluster 1, a total of 20 transcripts homologous to the genes encoding for photosynthesis were observed as up-regulated differentially expressed three days after harvest when all hardening accessions were analyzed together (Table 1), including chlorophyll a/b binding protein LHCB1 (8 transcripts), LHCA4 (2 transcripts) LHCB2 (2 transcripts), photosystem II protein psbX (2 transcripts). Those genes response to light stimulus and may be the triggers of this phenomenon. Three transcripts associated with cell wall organization were found encoding for fasciclin-type arabinogalactan protein, COB cellulose and glucan endo-1,3-beta-glucosidase. They are likely involved in the reinforcement of the cell wall (hardening). One transcript homologous to the gene related to the transcription factor TF-MYB was included in this group. However, it is important to note that genes involved in lipid metabolism namely lipase (3 transcripts) were found in this group.

**Table 1.** Candidate genes associated with post-harvest hardening in D. dumetorum tuber on 3DAH vs AH DEG on All accession 3DAH vs AH.

| Contig | LF2C | padj | Bin/KO | Gene\Name | Description |
| --- | --- | --- | --- | --- | --- |
| contig544.g2040 | 6.91 | 0.04740 | 21.4.1.1.3 | FLA | fasciclin-type AGP |
| contig278.g50 | 8.89 | 0.02720 | 21.1.2.2 | COB | regulatory protein |
| contig760.g29 | 18.35 | 0.03609 | 1.2.3/K05298 | GAPA | glyceraldehyde-3-phosphate dehydrogenase |
| contig119.g125 | 8.07 | 0.00170 | 1.1.6.1.1 | PGR5/PGRL1 | complex.component PGR5-like |
| contig678.g379 | 7.72 | 0.00000 | 1.1.4.1.4/K08910 | LHCA4 | chlorophyll a/b binding protein 4 |
| contig679.g24 | 7.98 | 0.00000 | 1.1.4.1.4/K08910 | LHCA4 | chlorophyll a/b binding protein 4 |
| contig549.g218 | 6.53 | 0.00000 | K02694 | psaF | photosystem I subunit III |
| contig222.g1555 | 5.27 | 0.00626 | 1.1.4.2.8/K02695 | psaH | photosystem I subunit VI |
| contig206.g10 | 5.55 | 0.00042 | 1.1.1.1.1/K08912 | LHCB1 | chlorophyll a/b binding protein 1 |
| contig206.g11 | 7.98 | 0.00000 | 1.1.1.1.1/K08912 | LHCB1 | chlorophyll a/b binding protein 1 |
| contig206.g6 | 7.12 | 0.00000 | 1.1.1.1.1/K08912 | LHCB1 | chlorophyll a/b binding protein 1 |
| contig206.g8 | 7.58 | 0.00000 | 1.1.1.1.1/K08912 | LHCB1 | chlorophyll a/b binding protein 1 |
| contig267.g402 | 5.81 | 0.00836 | 1.1.1.1.1/K08913 | LHCB2 | chlorophyll a/b binding protein 2 |
| contig355.g38 | 5.82 | 0.01516 | 1.1.1.1.1/K08913 | LHCB2 | chlorophyll a/b binding protein 2 |
| contig391.g20 | 6.24 | 0.00012 | 1.1.1.1.1/K08912 | LHCB1 | chlorophyll a/b binding protein 1 |
| contig391.g26 | 6.94 | 0.00000 | 1.1.1.1.1/K08912 | LHCB1 | chlorophyll a/b binding protein 1 |
| contig391.g28 | 5.72 | 0.00038 | 1.1.1.1.1/K08912 | LHCB1 | chlorophyll a/b binding protein 1 |
| contig391.g29 | 7.65 | 0.00000 | 1.1.1.1.1/K08912 | LHCB1 | chlorophyll a/b binding protein 1 |
| contig553.g402 | 4.31 | 0.04740 | 1.1.1.1.1/K08914 | LHCB3 | chlorophyll a/b binding protein 3 |
| contig565.g52 | 7.56 | 0.02366 | 1.1.1.1.1/K08912 | LHCB1 | chlorophyll a/b binding protein 1 |
| contig89.g1873 | 5.94 | 0.01452 | 1.1.1.6.2.1 | ELIP | LHC-related protein group.protein |
| contig544.g1881 | 5.17 | 0.04740 | 1.1.1.2.13 | 1.1.1.2.13/PsbX | PS-II complex.component |
| contig544.g1970 | 5.05 | 0.00905 | 1.1.1.2.13 | 1.1.1.2.13/PsbX | PS-II complex.component |
| contig267.g494 | 20.89 | 0.00000 | 15.5.2.1/K09422 | MYB | transcription factor |

In Fonkouankem (Table 2), 18 up-regulated genes encoding for cell wall organization including xylan O-acetyltranferase XOAT (5 transcripts), cellulose synthase CESA (3 transcripts), COB cellulose (2 transcripts) were found in cluster 2. The transcription factor MYB was the most abundant (4 transcripts) followed by DREB and NAC with 2 transcripts each of them. Photosynthesis genes LHCB1, LHCA4 were found with 2 transcripts each of them. However, genes encoding for phenolic metabolism were enriched with 2 genes cinnamate 4-hydroxylase (2 transcripts) and phenylalanine ammonia lyase (2 transcripts). Likewise, lipase (3 transcripts) was recorded in this group.

**Table 2.**
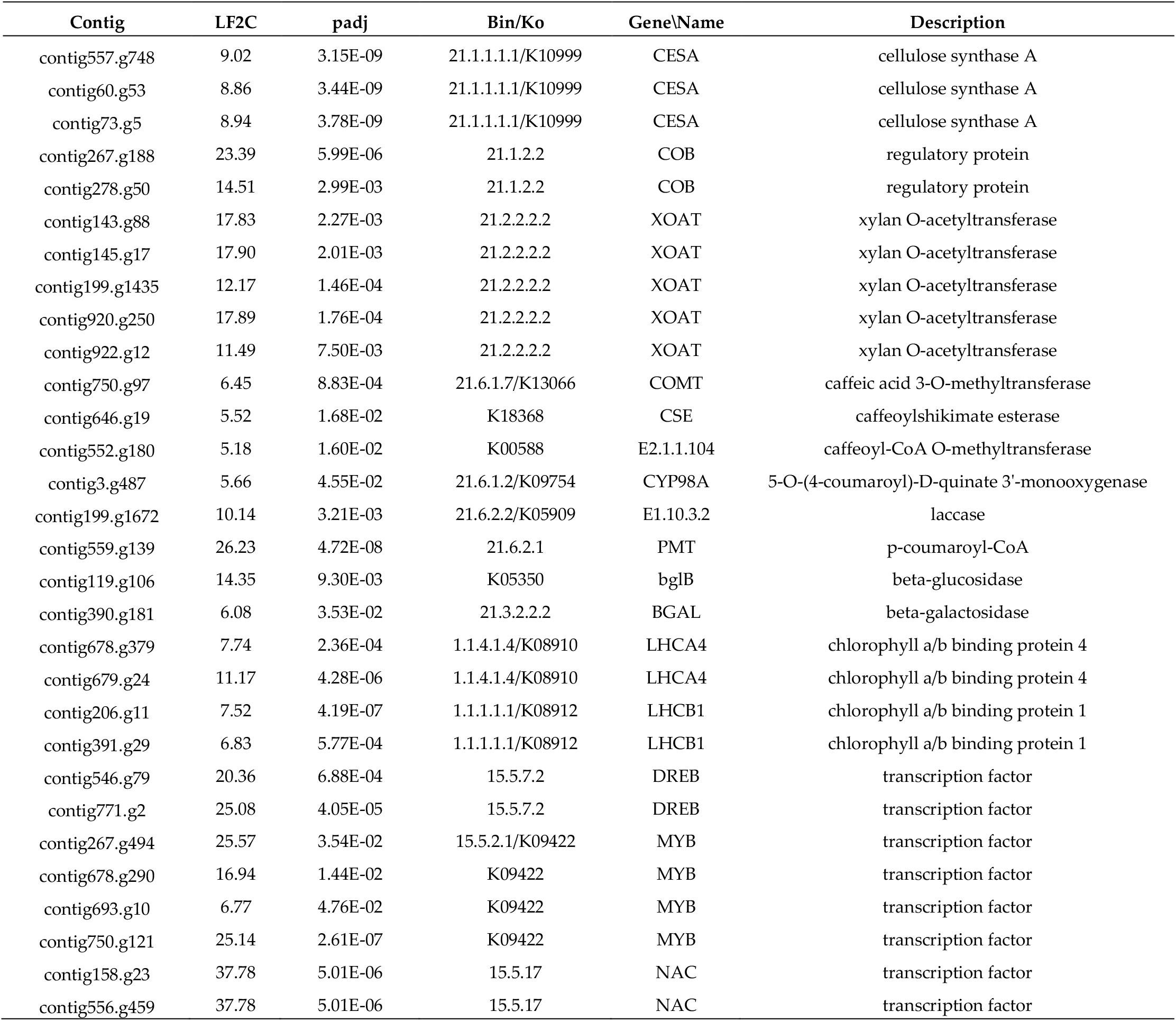
Candidate genes associated with post-harvest hardening in D. dumetorum tuber on Fonkouankem 3DAH vs AH.

| Contig | LF2C | padj | Bin/Ko | Gene\Name | Description |
| --- | --- | --- | --- | --- | --- |
| contig557.g748 | 9.02 | 3.15E-09 | 21.1.1.1/K10999 | CESA | cellulose synthase A |
| contig60.g53 | 8.86 | 3.44E-09 | 21.1.1.1/K10999 | CESA | cellulose synthase A |
| contig73.g5 | 8.94 | 3.78E-09 | 21.1.1.1/K10999 | CESA | cellulose synthase A |
| contig267.g188 | 23.39 | 5.99E-06 | 21.1.2.2 | COB | regulatory protein |
| contig278.g50 | 14.51 | 2.99E-03 | 21.1.2.2 | COB | regulatory protein |
| contig143.g88 | 17.83 | 2.27E-03 | 21.2.2.2 | XOAT | xylan O-acetyltransferase |
| contig145.g17 | 17.90 | 2.01E-03 | 21.2.2.2 | XOAT | xylan O-acetyltransferase |
| contig199.g1435 | 12.17 | 1.46E-04 | 21.2.2.2 | XOAT | xylan O-acetyltransferase |
| contig920.g250 | 17.89 | 1.76E-04 | 21.2.2.2 | XOAT | xylan O-acetyltransferase |
| contig922.g12 | 11.49 | 7.50E-03 | 21.2.2.2 | XOAT | xylan O-acetyltransferase |
| contig750.g97 | 6.45 | 8.83E-04 | 21.6.1.7/K13066 | COMT | caffeic acid 3-O-methyltransferase |
| contig646.g19 | 5.52 | 1.68E-02 | K18368 | CSE | caffeoylshikimate esterase |
| contig552.g180 | 5.18 | 1.60E-02 | K00588 | E2.1.1.104 | caffeoyl-CoA O-methyltransferase |
| contig3.g487 | 5.66 | 4.55E-02 | 21.6.1.2/K09754 | CYP98A | 5-O-(4-coumaroyl)-D-quinic acid 3'-monooxygenase |
| contig199.g1672 | 10.14 | 3.21E-03 | 21.6.2.2/K05909 | E1.10.3.2 | laccase |
| contig559.g139 | 26.23 | 4.72E-08 | 21.6.2.1 | PMT | p-coumaroyl-CoA |
| contig119.g106 | 14.35 | 9.30E-03 | K05350 | bglB | beta-glucosidase |
| contig390.g181 | 6.08 | 3.53E-02 | 21.3.2.2 | BGAL | beta-galactosidase |
| contig678.g379 | 7.74 | 2.36E-04 | 1.1.4.1.4/K08910 | LHCA4 | chlorophyll a/b binding protein 4 |
| contig679.g24 | 11.17 | 4.28E-06 | 1.1.4.1.4/K08910 | LHCA4 | chlorophyll a/b binding protein 4 |
| contig206.g11 | 7.52 | 4.19E-07 | 1.1.1.1.1/K08912 | LHCB1 | chlorophyll a/b binding protein 1 |
| contig391.g29 | 6.83 | 5.77E-04 | 1.1.1.1.1/K08912 | LHCB1 | chlorophyll a/b binding protein 1 |
| contig546.g79 | 20.36 | 6.88E-04 | 15.5.7.2 | DREB | transcription factor |
| contig771.g2 | 25.08 | 4.05E-05 | 15.5.7.2 | DREB | transcription factor |
| contig267.g494 | 25.57 | 3.54E-02 | 15.5.2.1/K09422 | MYB | transcription factor |
| contig678.g290 | 16.94 | 1.44E-02 | K09422 | MYB | transcription factor |
| contig693.g10 | 6.77 | 4.76E-02 | K09422 | MYB | transcription factor |
| contig750.g121 | 25.14 | 2.61E-07 | K09422 | MYB | transcription factor |
| contig158.g23 | 37.78 | 5.01E-06 | 15.5.17 | NAC | transcription factor |
| contig556.g459 | 37.78 | 5.01E-06 | 15.5.17 | NAC | transcription factor |

In all hardening accession and the combining analysis of all three accessions together, annotation with several MYB database identified putative MYB genes (MYB54, MYB52, MYB73, MYB70, MYB44, MYB77, MYB46, MYB83, MYB9, MYB107, MYB93, MYB53, MYB92) associated with cell wall modifications (Supplementary S4).

### 2.5. Comprehensive difference between harden and non-harden accessions

Pairwise comparisons of accessions that do harden to the accession that does not harden in different stage after harvest showed that up-regulated genes were enriched mostly in cellular process, cellular anatomical entity and intracellular 3DAH and 14DAH (Figure 7, Supplementary S5). Besides, KEGG enrichment revealed that metabolic pathways were the most enriched with 10, 8 and 5 up-regulated genes 3DAH for Bayangam vs Ibo, Fonkouankem vs Ibo and Bangou vs Ibo respectively (Figure 7 A, B, C). This pathway was followed by biosynthesis of secondary metabolites with 6, 5, and 5 up-regulated genes for Bayangam 2 vs Ibo, Fonkouankem 1 vs Ibo sweet 3 and Bangou 1 vs Ibo sweet 3 respectively. Those pathways were the most enriched as well 14DAH (Supplementary S6). MapMan annotation showed that cell wall organization was predominantly enriched when comparing Bangou 1 to Ibo sweet 3 and Fonkouankem 1 vs Ibo sweet 3 3DAH. Whereas protein modification was particularly enriched for Bayangam 2 vs Ibo sweet 3. However, cell organization, protein modification and RNA biosynthesis belong to the top 7 of the most enriched pathways 3DAH. On the contrary, protein modification was the most enriched irrespective of the comparison 14DAH (Supplementary S6). Venn diagram of the annotation revealed 5 common up-regulated genes potentially involved in the hardening process among the accessions that do harden comparing to the non-hardening accession Ibo sweet 3. Those genes encoding for chalcone synthase, diterpene synthase, transcription factor MYB, xylan O-acetyltransferase (XOAT), lignin laccase (Figure 7 D).

**Figure 7.**
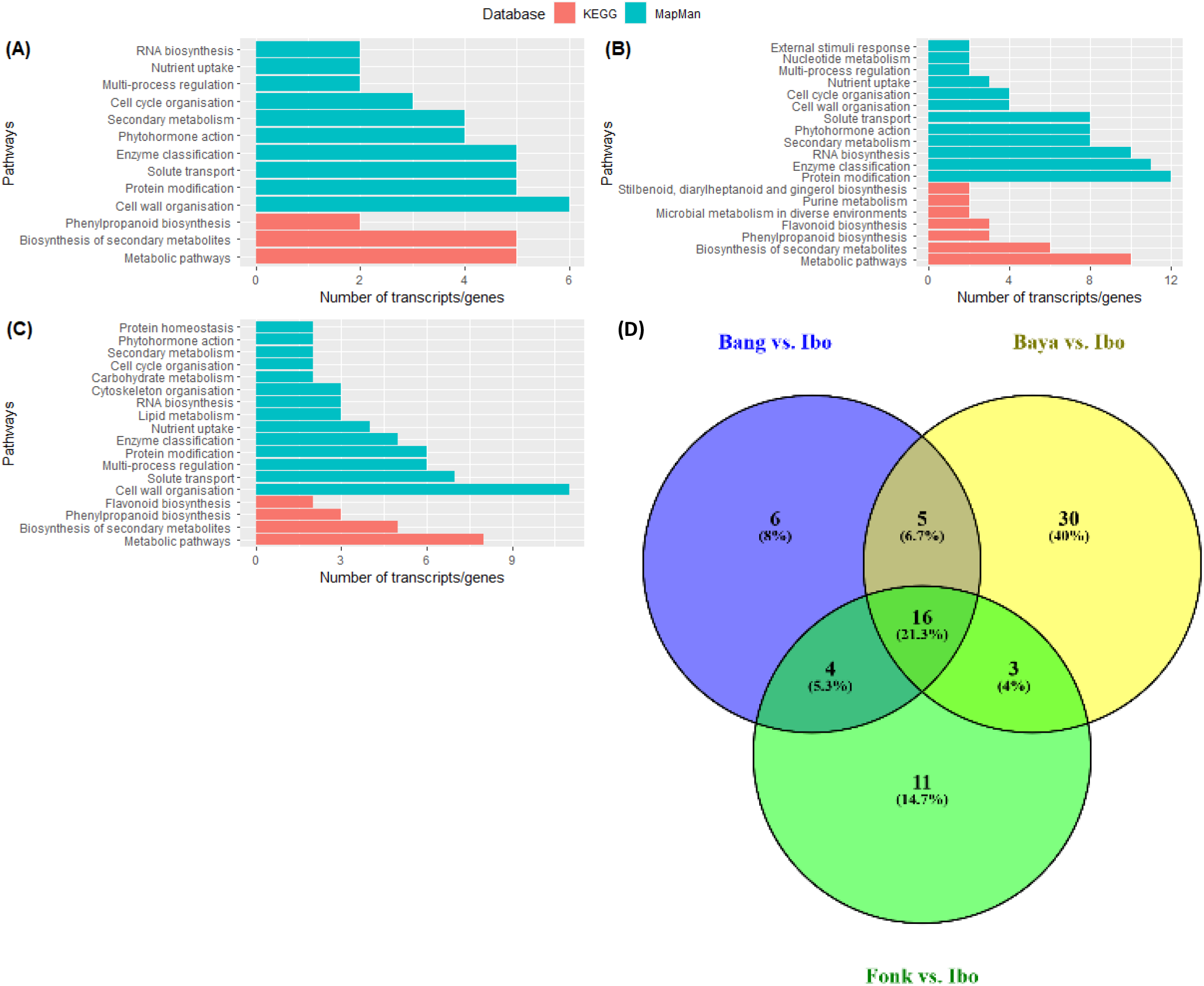
Functional classification of up-regulated DEG 3DAH based on the comparison of hardened accessions against the non-hardening accessions. (A), (B) and (C) the most enriched pathways 3 DAH on Bangou 1 vs. Ibo sweet 3, Bayangam 2 vs. Ibo sweet 3 and Fonkouankem 1 vs. Ibo sweet 3 respectively. (D) venn diagram of the most enriched pathways on Bangou 1 vs. Ibo sweet 3, Bayangam 2 vs. Ibo sweet 3 and Fonkouankem 1 vs. Ibo sweet 3. Green represents pathway annotation with MapMan database, and red represents pathway annotation with KEGG database.

**Figure 7.**
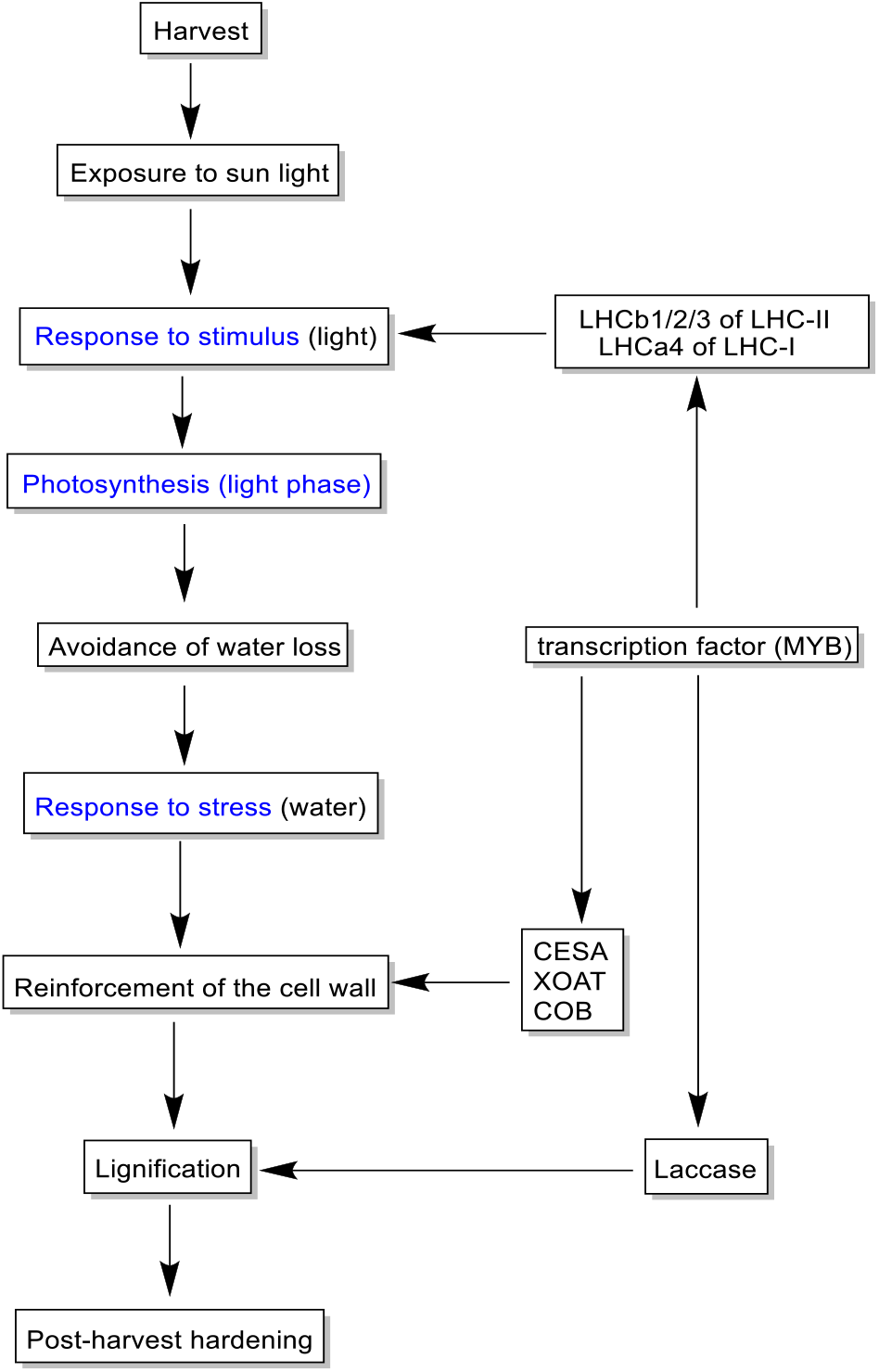
Putative mechanism of the PHH on *D. dumetorum*. Blue represents GO annotation.

## 3. Discussion

The post-harvest hardening of *D. dumetorum* tubers has been extensively studied. regarding the biochemical and physical aspects [1]-17]. Based on our study, we reported genes that differential expressed and up-regulated AH. This demonstrates that the PHH on *D. dumetorum* Tuber is likely controlled by genes. Our results showed that the number of the up-regulated genes was abundant 3DAH and then decreased 14DAH. This suggest that the PHH predominantly occurs few days after harvest. This is consistent with previous studies [1,8,18] showing a substantial increase of the hardness the first 3 DAH.

Functional analysis via KEGG enrichment revealed that most genes were involved in pathways of secondary metabolites. These genes were involved in photosynthesis, RNA biosynthesis (transcription factors), cell wall organisation. In order to understand causes of this phenomenon, GO enrichment revealed that many genes were involved in cellular process, response to stimulus and metabolic process, response to stress. These results prove that the PHH on *D. dumetorum* is a cellular and metabolic process in response to stimulus leading to stress.

Indeed, [1] reported that the PHH on *D. dumetorum* is associated with an increase in sugar and structural polysaccharides (cellulose, hemicellulose, and lignin). Later, [18] associated it with a decrease of phytate and total phenols. However, these authors failed to address causes of this phenomenon. Cellular processes are triggered by stimulus, an investigation of genes related to response to stimulus revealed photosynthetic genes LHCB1,2,3 and LCH4 were up regulated 3DAH. Those genes are light-harvesting chlorophyll a/b binding antenna responsible for photons capture. This suggests that *D. dumetorum* tubers are capable of photosynthesis. In the field, *D. dumetorum* tubers turn green under the yam skin (on the surface) were they are exposed to sun light (Supplementary S7). Unlike potatoes, greening occurs only in the field but not in storage. Photosynthesis implies that the sunlight energy capture through photons is used to extract electron from water leading to the synthesis of adenosine triphosphate ATP and nicotinamide adenine dinucleotide phosphate NADPH [19], highlighting the importance of water in this process. After harvest, tubers are exposed to the external environment with no possibility of water absorption. This likely leads to a stress process as revealed by GO term analysis in relation with water limitation. In fact, a rapid decrease of water on tuber after harvest was reported [1–18], probably due in majority to this putative photosynthetic activity of *D. dumetorum* tubers. Thus, the PHH of *D. dumetorum* tubers appears as a mechanism to limit water loss.

Mechanism of limitation of water loss in plant has been extensively associated with the reinforcement of the cell wall [20]. Indeed, [18] reported a decrease of water absorption by tubers after harvest suggesting that the cell wall permeability decreases during the storage. Genes related to cell wall organisation xylan O-acetyltranferase XOAT, cellulose synthase CESA, corncob cellulose COB cellulose were predominantly up-related after harvest. This confirms biochemical changes associated with the PHH of *D. dumetorum* tubers [1]-18]. They observed an increase in various cell wall polysaccharide such as cellulose, hemicellulose and lignin during storage. Cellulose synthase encodes for cellulose biosynthesis [21] and COB regulate the orientation of cellulose microfibrils whereas xylan O-acetyltranferase XOAT encode for hemicellulose (xylan) [22]. These cell wall polysaccharides play an important role as a protective barrier in response to various environmental perturbations. Accumulation and deposition of these polysaccharides inside primary walls reinforces the strength and rigidity of the cell wall and are probably a key component of the plant response to environment factors [20]. It suggests that cellulose and lignin are key cell wall polymers responsible for cell wall rigidification during the PHH on *D. dumetorum*.

Many biological processes are controlled by the regulation of gene expression at the level of transcription. Transcription factors TFs are key players in controlling cellular processes. Among those TFs, MYB family is large and involved in controlling diverse processes such as responses to abiotic and biotic stresses [23]. Our results showed that TF from MYB family was predominantly expressed and up regulated after harvest. This result suggests that transcription factors from MYB family may be potentially involved in the mechanism of post-harvest hardening. [24] demonstrated the role of an MYB TF family in response to water stress from stem of a plant tree birch through lignin deposition, secondary cell wall thickness and the expression of genes in secondary cell wall formation.

Pairwise comparison of the hardened accessions and the non-hardened accession confirmed that the PHH is a cellular and metabolic process leading to the cell wall modification. However, it is interesting to note that protein modifications seem to occur predominantly after hardness from 3 to 14 DAH. This could explain the poor sensory qualities of hardened tubers such as coarseness in the mouth [25]. Five common genes were found up-regulated in the hardened accessions and down-regulated in Ibosweet 3 3DAH. Those genes are chalcone synthase, diterpene synthase, transcription factor MYB, xylan O-acetyltransferase and lignin laccase. Chalcone synthase is a key enzyme of the flavonoids/isoflavonoid biosynthesis pathway and is induced in plants under stress conditions [26]. Laccase catalyse the oxidation of phenolic substrates using oxygen as electron acceptor. Laccase has been recognized in the lignification process through the oxidation of lignin precursors. Indeed, [27] demonstrated an involvement of laccase genes in lignification as response to adaptation to abiotic stresses in *Eucalytus*.

Based on our results, the PHH seems to be governed by differentially expressed genes in a metabolic network, which is attributed to the exposure to external environment or sun light. Therefore, a putative model of the hardening mechanism and the regulatory network associated was proposed (Figure 8). After harvest, yam tubers are exposed to the external environment particularly to sun light. This environmental factor acts as the first signal to stimulate photosynthetic genes involved in photons capture namely LHCB1, LHCB2, LHCB3 and LHCA4. The absorption of photons implies loss of electrons which is replaced by electrons from the spitting water through photolysis [28]. This activity implies the necessity of a continued electron supply through the breakdown of water molecule. However, tubers are detached from roots with no possibility of water absorption. Therefore, a signal is given to reinforce the cell wall in order to avoid loss of water from the tubers via the up regulation of CESA, XOAT and COB genes. This reinforcement of the cell wall implies firstly, an accumulation of cell wall polysaccharide such as cellulose hemicellulose during the first days of storage. Secondly, probably from the third day after harvest starts the lignification process controlling laccase genes. This overall process is likely controlled transcription factor MYB.

## 4. Materials and Methods

### 4.1. Plant materials

Four accessions have been collected from various localities in the main growing regions of yam (West and South-West) in Cameroon and one from Nigeria based on the analysis of [9]. These accessions were planted in pots in the greenhouse of the botanic garden of the University of Oldenburg under controlled conditions at 25 °C. They are available upon request.

### 4.2. Sample preparation

Three tubers of each accession were randomly collected 4 months after emergence (ME), 9 ME (Harvest time AH), 3 days after harvest (3DAH) and 14 DAH. Collected tubers were washed and their skin peeled off. Then, the samples will be immediately frozen in liquid nitrogen and stored at – 80 °C prior to RNA isolation.

### 4.3. RNA-Seq extraction

The stored tubers (– 80 °C) were immediately lyophilized. Total RNA was extracted from 48 samples using innuPREP Plant RNA Kit (Analytik Jena AG, Germany). The RNA quality was analysed using a spectrophotometer (Nano-Drop Technologies, Wilmington, DE, USA). RNA Integrity Number (RIN) values were determined using a Bioanalyzer 2100 (Agilent Technologies, Santa Clara, CA) to ensure all samples had a RNA integrity number (RIN) above 6.

### 4.4. Library construction and Illumina sequencing

We constructed cDNA libraries comprising 48 RNA samples using the Universal Plus mRNA-Seq offered by NuQuant (Tecan Genomics, Inc California, USA). Paired-end (2 × 150 bp) sequencing of the cDNA libraries was performed on the Illumina HiSeq 2000 (Illumina Inc., San Diego, CA, USA).

### 4.5. Data processing and functional analysis

Low quality reads were filtered using TrimGalore v 0.6.5 (https://github.com/FelixKrueger/TrimGalore/releases) with the following parameters --length 36 -q 5 --stringency 1 –e 0.1. The filtered reads were aligned to the reference genome of *D. dumetorum* [10] with STAR v 2.7.3a [29] with default parameters. The aligned reads in BAM files were sorted and indexed using SAMtools v 1.9 [30]. The number of reads that can be assigned uniquely to genomic features were counted using the function SummarizeOverlaps of the R package GenomicAlignments v1.20.1 [31] with mode=“Union”, singleEnd=FALSE, ignore.strand=TRUE, fragments=TRUE as parameters.

Two programs DESeq2 [32] and edgeR [33] were deployed to analyze differentially expressed genes (DEGs) between conditions and the interaction conditions x accessions. Gene with p-adjusted value < 0.05 and log2 fold change > 2 were considered as significantly expressed genes. False discovery rate FDR threshold was < 0.05. We performed a basic time course experiment to assess genes that change their expression after harvest using Deseq2 [32]. Metabolic pathway assignments of DEGs were based on the KEGG Orthology database using the KAAS system [34]. The final pathway analyses were mostly based on the tool Mercator4 and Mapman4 [35]. In addition, differential expressed MYB genes were functional annotated based on several datasets *Arabidopsis thaliana* MYBs [36], *Beta vulgaris* MYBs [37], *Musa acuminata* MYBs [38], *Croton tiglium* MYBs [39], *Dioscorea rotundata* MYBs and *Dioscorea dumetorum* MYBs via KIPEs (https://github.com/bpucker/KIPEs). GO term assignment and enrichment were performed using Blast2GO [40] via OmicsBox with cutoff 55, Go weight 5, e-value 1.e-6, HSP-hit coverage cutoff 80 and hit filter 500. Co-expression analysis was carried out using k-means method and the number of cluster was determined through the sum of squared error and the average silhouette width.

## 5. Conclusions

In this study, for the first time differentially expressed genes after harvest and during yam storage was investigated through RNA-Seq. The evidence from this study suggests that the PHH on *D. dumetorum* is a cellular and metabolic process involving a combined action of several genes as response to environmental stress due to sun and water. Genes encoding for cell wall polysaccharide constituents were found significantly up-regulated suggesting that they directly responsible for the hardness of *D. dumetorum* tubers. It is worth noticing that many genes encoding for light-harvesting chlorophyll a/b binding proteins were as well significantly up regulated after harvest. This support the idea that sunlight is the trigger element of the PHH manifested by the strengthen of cell call in order to avoid water loss useful for a putative photosynthesis activity. These findings add substantially to our understanding of hardening on *D. dumetorum* and provide the framework for molecular breeding against the PHH on *D. dumetorum*.

## Supplementary Materials

Supplementary S1: Statistic of clean reads mapped to *D. dumetorum* reference genome, Supplementary S2: Number of DEGs based on the combined analysis of the three hardening accessions 4MAE and after harvest, Supplementary S3: Group resulting from Cluster analysis of DEGs 3DAH among the different sampling times for Bangou, Bayangam, Fonkouankem, and the combined analysis of the three hardening accessions, Supplementary S4: Phylogenetic tree of candidate MYB genes in Bangou, Bayangam, Fonkouankem, and the combined analysis of the three hardening accessions, Supplementary S5: GO enrichment of up-regulated DEG 3DAH and 14DAH based on the comparison of hardening accessions against the non-hardening accession, Supplementary S6: Functional classification of up-regulated DEG 14DAH based on the comparison of hardening accessions against the non-hardening accession. (A), (B) and (C) the most enriched pathways 14 DAH in Bangou 1 vs. Ibo sweet 3, Bayangam 2 vs. Ibo sweet 3, and Fonkouankem 1 vs. Ibo sweet 3, respectively. Green bars represent pathway annotation with the MapMan database, and red bars represent pathway annotation with the KEGG database, Supplementary S7: Greening of young *D. dumetorum* tuber exposed to sunlight as opposed to the non-greening one.

## Author Contributions

Conceptualization, C.S. and D.C.A.; methodology, C.S., E.M., and S.L.; software, C.S.; validation, C.S.; formal analysis, C.S.; investigation, C.S., E.M.; resources, C.S.; data curation, C.S.; writing—original draft preparation, C.S.; writing—review and editing, D.C.A., S.L.; visualization, C.S.; supervision, D.C.A., S.L.; project administration, D.C.A., E.M.; funding acquisition, DCA., C.S. All authors have read and agreed to the published version of the manuscript.

## Funding

This research was funded by Alexander von Humboldt-Stiftung, grant number 1128007-NGA-IP and by Deutscher Akademischer Austauschdienst DAAD, grant number 57299294.

## Data Availability Statement

Data are available upon request.

## Acknowledgments

We would like to thank Dr. Boas Pucker for helping in the annotation with KIPEs.

## Conflicts of Interest

The authors declare no conflict of interest.

## Notes

### Competing Interest Statement

The authors have declared no competing interest.

