## Supplementary S1 for "Transcriptome sequence reveals candidate genes involving in the post-harvest hardening of trifoliate yam *Dioscorea dumetorum*"

| Accessions | Conditions | Total cleaned reads | Total (%) | Uniquely mapped reads (%) | Total (%) |
|---|---|---|---|---|---|
| Bangou 1 | 4MAE | 17893940 | 56.61 | 36.51 | 45.13 |
| Bangou 1 | 4MAE | 18949046 | 93.49 | 60.09 | 69.42 |
| Bangou 1 | 4MAE | 20612907 | 92.51 | 58.2 | 69.05 |
| Bangou 1 | AH | 21284551 | 96.13 | 59 | 74.38 |
| Bangou 1 | AH | 20481906 | 91.77 | 53.96 | 70.26 |
| Bangou 1 | AH | 17929078 | 93.36 | 53.29 | 78.56 |
| Bangou 1 | 3DAH | 19902140 | 95.5 | 56.66 | 74 |
| Bangou 1 | 3DAH | 24032161 | 93.56 | 51.76 | 78.23 |
| Bangou 1 | 3DAH | 14752529 | 86.88 | 52.2 | 70.09 |
| Bangou 1 | 14DAH | 18279097 | 77.73 | 45.16 | 63.81 |
| Bangou 1 | 14DAH | 20344788 | 91.64 | 56.5 | 74.26 |
| Bangou 1 | 14DAH | 4536123 | 93.3 | 58.73 | 69.95 |
| Bayangam 2 | 4MAE | 53929557 | 96.63 | 63.53 | 71.97 |
| Bayangam 2 | 4MAE | 22569645 | 91.81 | 60.56 | 71.45 |
| Bayangam 2 | 4MAE | 24937580 | 88.9 | 57.83 | 65.31 |
| Bayangam 2 | AH | 21708627 | 82.96 | 52.63 | 61.75 |
| Bayangam 2 | AH | 23997962 | 97.29 | 53.7 | 80.6 |
| Bayangam 2 | AH | 31335502 | 94.09 | 49.7 | 78.78 |
| Bayangam 2 | 3DAH | 24284434 | 94.56 | 54.66 | 72.63 |
| Bayangam 2 | 3DAH | 22565017 | 93.62 | 56.11 | 76.8 |
| Bayangam 2 | 3DAH | 20803207 | 96.21 | 58.49 | 74.35 |
| Bayangam 2 | 14DAH | 27593312 | 90.11 | 55.12 | 72.04 |
| Bayangam 2 | 14DAH | 18519729 | 95.9 | 62.63 | 71.94 |
| Bayangam 2 | 14DAH | 19713286 | 96 | 60.25 | 70.93 |
| Fonkouankem 1 | 4MAE | 22166474 | 94.25 | 62.53 | 71.25 |
| Fonkouankem 1 | 4MAE | 12111303 | 84.48 | 55.59 | 61.48 |
| Fonkouankem 1 | 4MAE | 20286805 | 85.76 | 56.75 | 61.04 |
| Fonkouankem 1 | AH | 13126435 | 93.29 | 53.1 | 74.92 |
| Fonkouankem 1 | AH | 19560305 | 92.51 | 55.3 | 73.54 |
| Fonkouankem 1 | AH | 13307694 | 93.43 | 56 | 70.66 |
| Fonkouankem 1 | 3DAH | 22500916 | 92.21 | 56.99 | 71.71 |
| Fonkouankem 1 | 3DAH | 17371457 | 96.19 | 56.96 | 74.36 |
| Fonkouankem 1 | 3DAH | 11617175 | 95.44 | 57.81 | 67.16 |
| Fonkouankem 1 | 14DAH | 20060449 | 93.44 | 62.04 | 68.06 |
| Fonkouankem 1 | 14DAH | 15829240 | 86.93 | 56.82 | 71.64 |
| Fonkouankem 1 | 14DAH | 15279090 | 86.98 | 56.88 | 63.02 |
| Ibosweet 3 | 4MAE | 16672162 | 93.04 | 59.72 | 68.71 |
| Ibosweet 3 | 4MAE | 13924548 | 91.2 | 58.68 | 68.4 |
| Ibosweet 3 | 4MAE | 30873744 | 91.99 | 57.53 | 70.07 |
| Ibosweet 3 | AH | 7881301 | 93.36 | 55.37 | 73.71 |
| Ibosweet 3 | AH | 14683717 | 94.95 | 56.98 | 75.31 |
| Ibosweet 3 | AH | 14787799 | 95.71 | 61.83 | 72.2 |
| Ibosweet 3 | 3DAH | 14413912 | 95.55 | 59.9 | 73.23 |
| Ibosweet 3 | 3DAH | 21994748 | 94.17 | 56.54 | 73.06 |
| Ibosweet 3 | 3DAH | 17962629 | 95.22 | 55.32 | 73.62 |
| Ibosweet 3 | 14DAH | 15590855 | 91.61 | 58.23 | 68.25 |
| Ibosweet 3 | 14DAH | 21780220 | 84.31 | 52.77 | 65.4 |
| Ibosweet 3 | 14DAH | 18583946 | 95.04 | 60.53 | 72.64 |
