## Supplementary figures and images for "Transcriptome sequence reveals candidate genes involving in the post-harvest hardening of trifoliate yam *Dioscorea dumetorum*"

### Supplementary S2

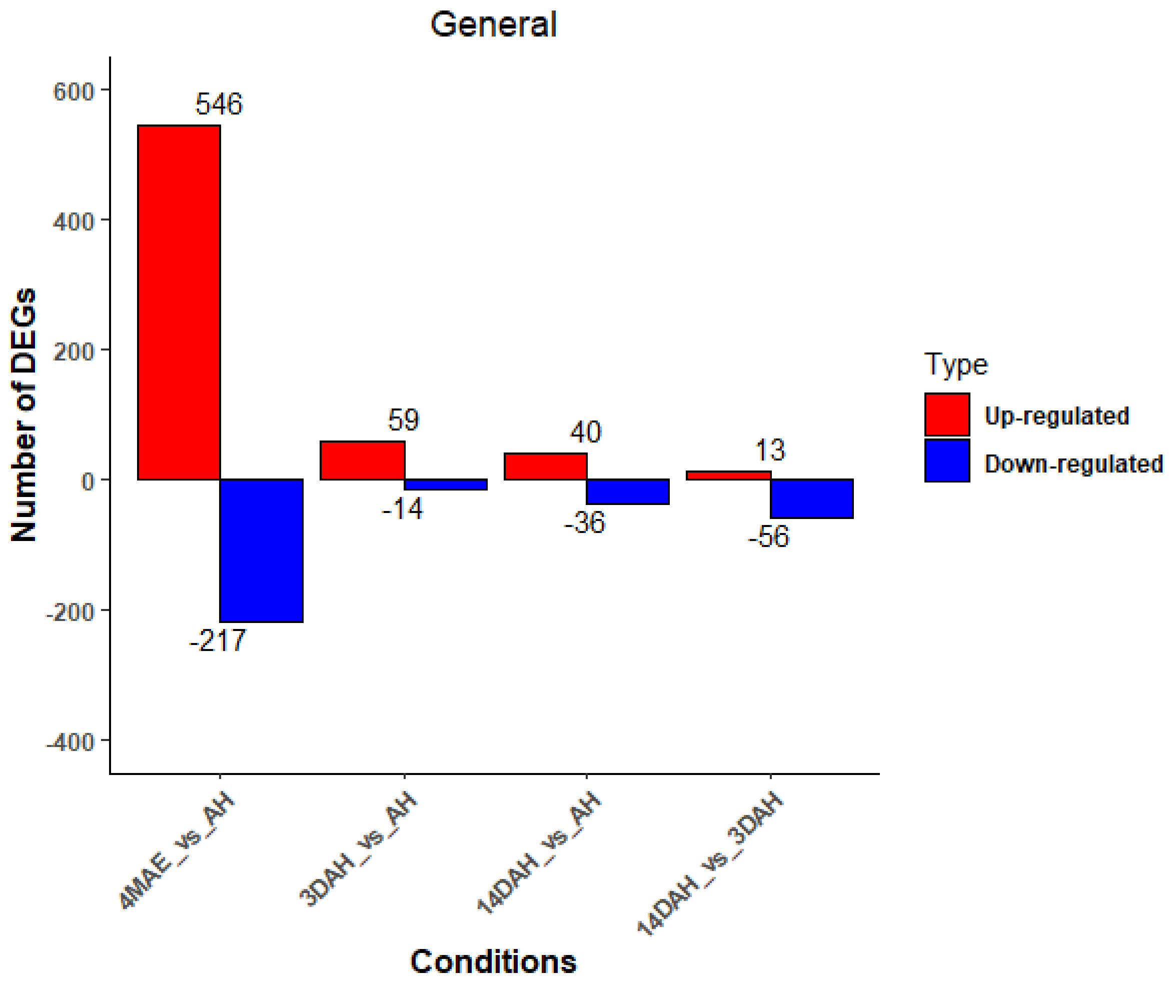

### Supplementary S6

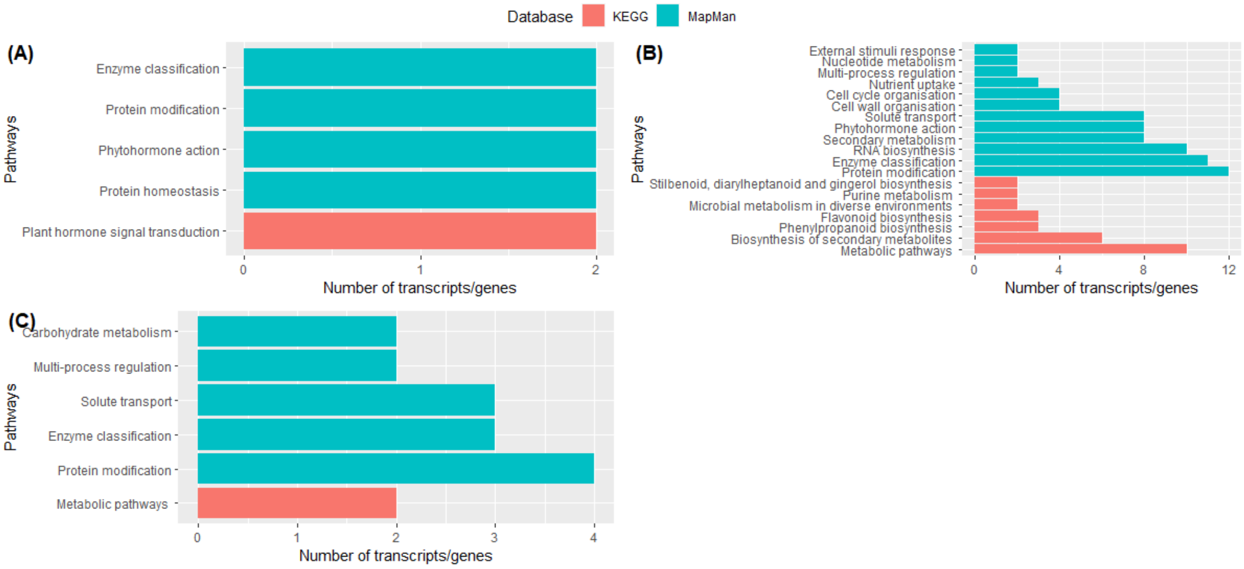

### Supplementary S7

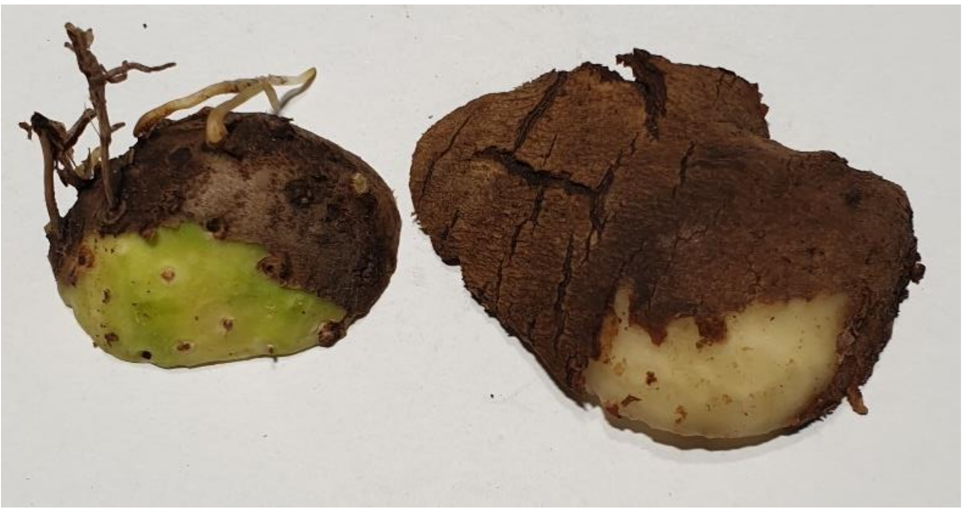
