## Supplementary S5 for "Transcriptome sequence reveals candidate genes involving in the post-harvest hardening of trifoliate yam *Dioscorea dumetorum*"

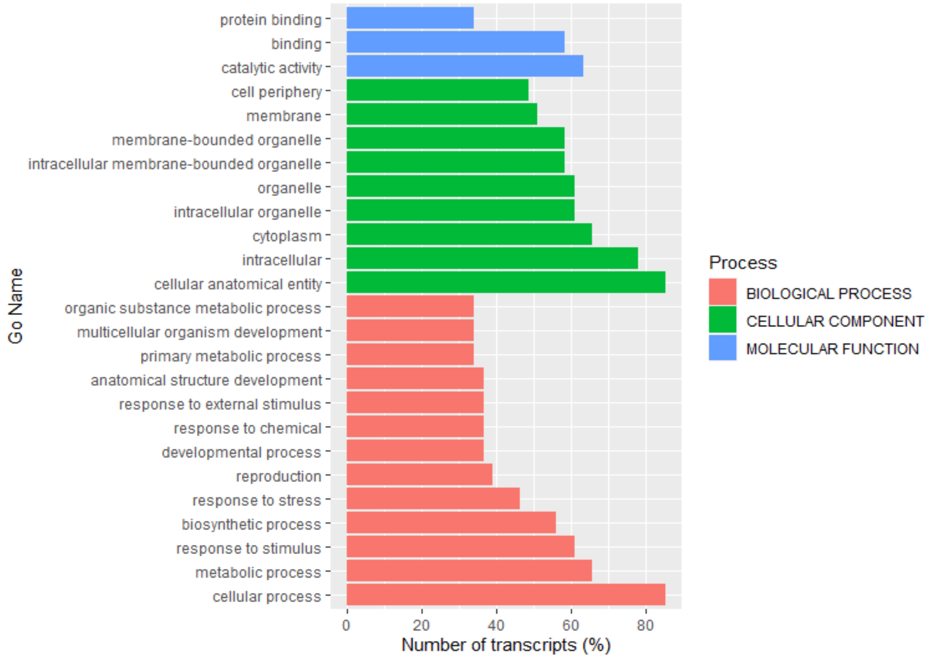


Bangou 1 vs. Ibo sweet 3 3DAHvsAH

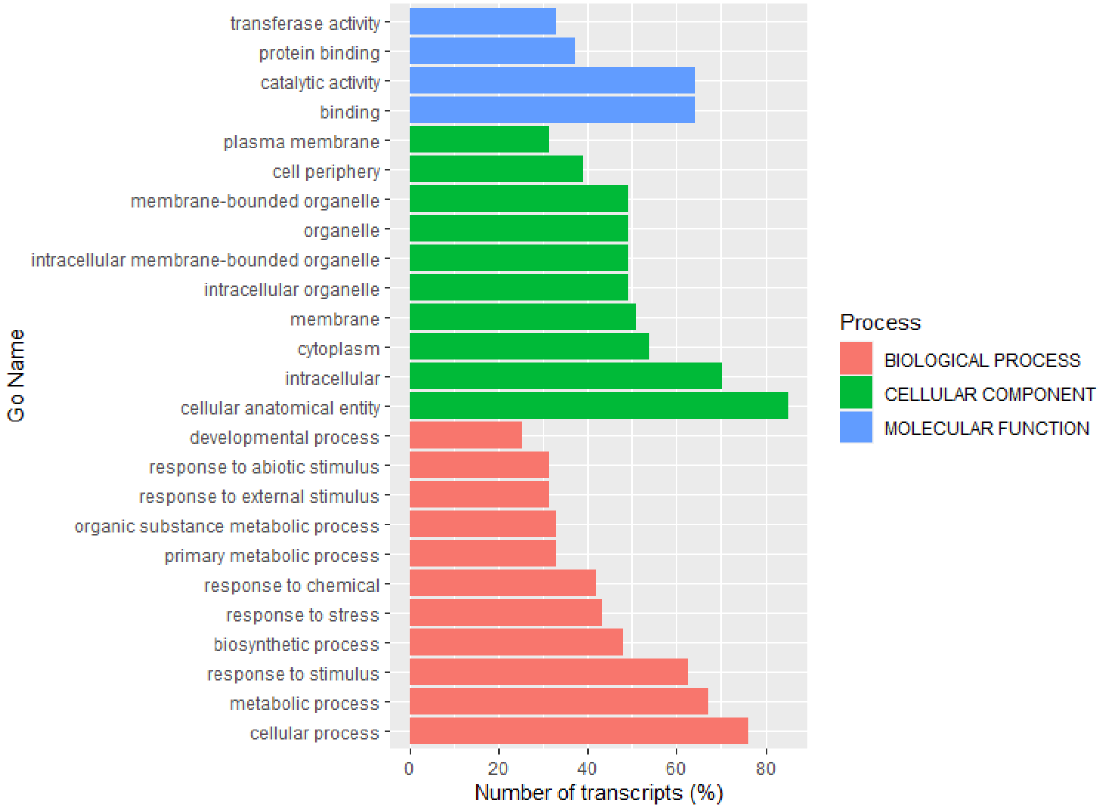


Bayangam 2 vs. Ibo sweet 3 3DAHvsAH

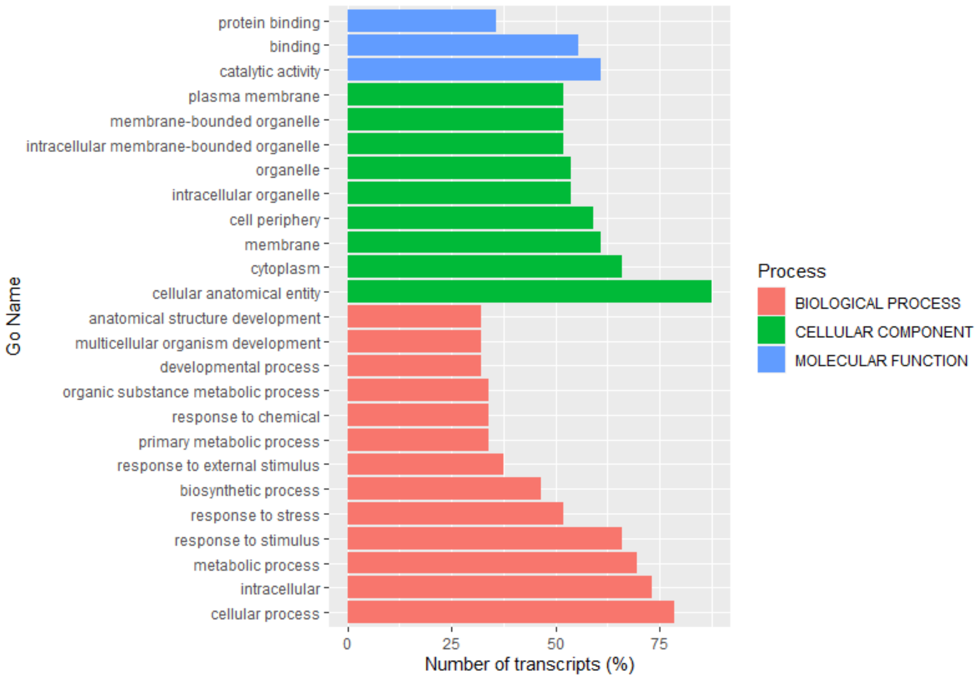


Fonkouankem 1 vs. Ibo sweet 3 3DAHvsAH

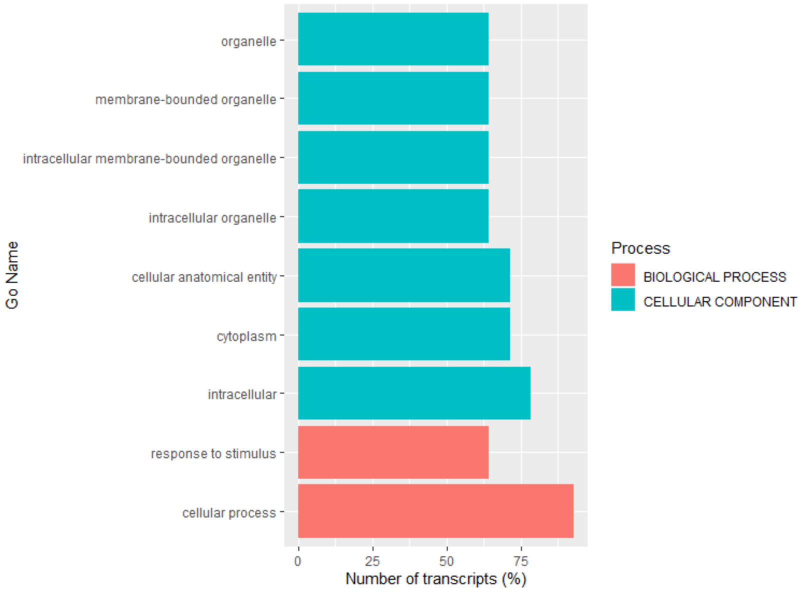


Bangou 1 vs. Ibo sweet 3 14DAHvsAH

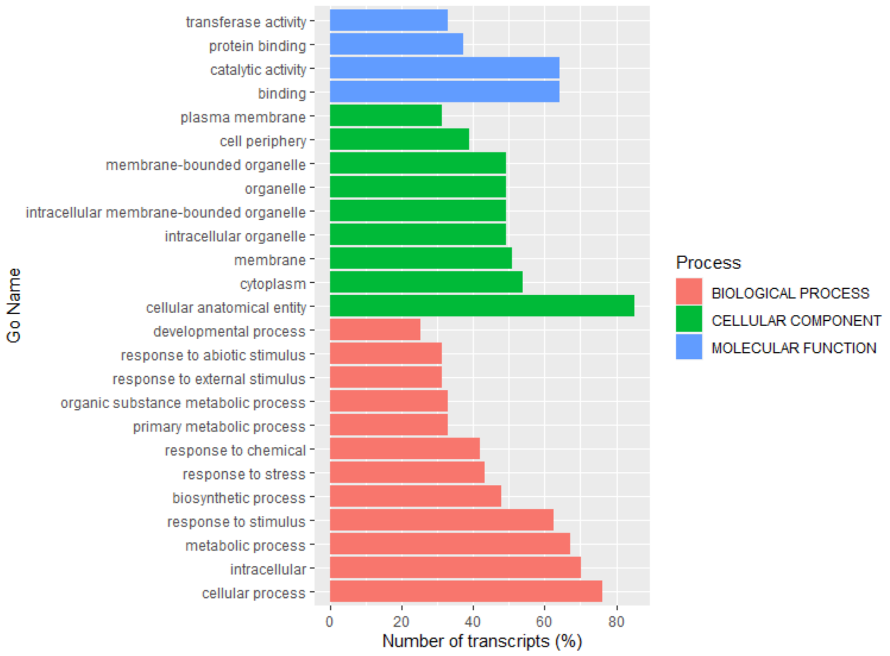


Bayangam 2 vs. Ibo sweet 3 14DAHvsAH

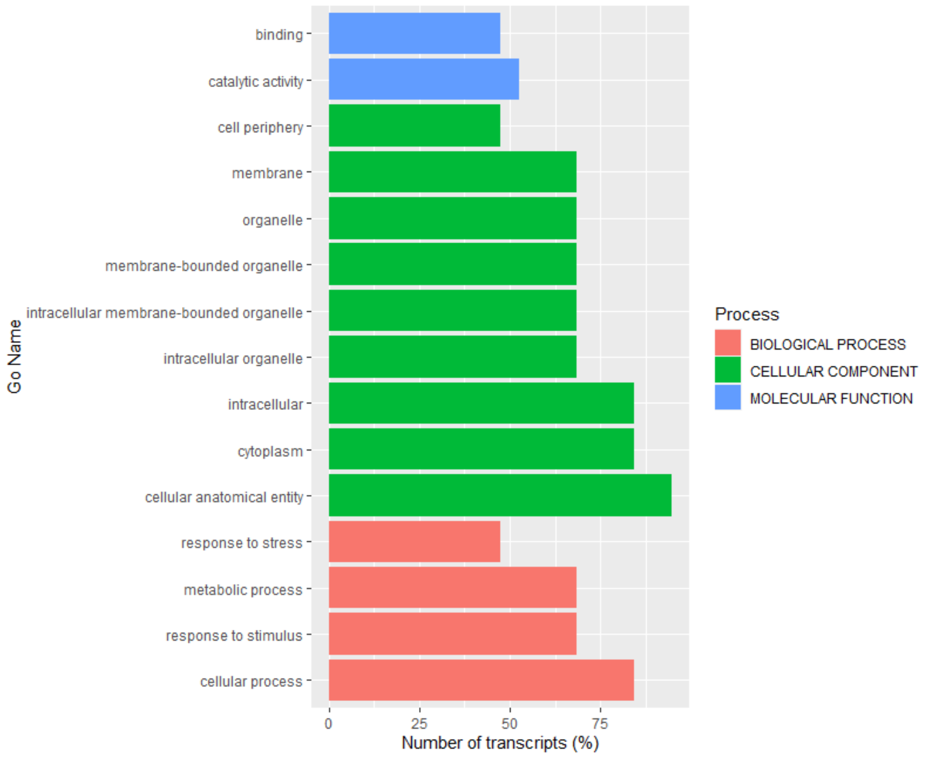


Fonkouankem 1 vs. Ibo sweet 3 14DAHvsAH
